# Ampk activation by glycogen expenditure primes the exit of naïve pluripotency

**DOI:** 10.1101/2023.05.19.541467

**Authors:** Seong-Min Kim, Eun-Ji Kwon, Ji-Young Oh, Han Sun Kim, Sunghyouk Park, Goo Jang, Jeong Tae Do, Keun-Tae Kim, Hyuk-Jin Cha

## Abstract

Embryonic and epiblast stem cells in pre-and post-implantation embryos are characterized by their naïve and primed states, respectively, which represent distinct phases of pluripotency. Thus, the cellular transition from naïve to primed pluripotency recapitulates a drastic metabolic and cellular remodeling after implantation to adapt to changes in extracellular conditions. Here, we found that inhibition of Ampk occurred during naïve transition with two conventional inhibitors (2i) of the Mek1 and Gsk3β pathways. The accumulation of glycogen due to the inhibition of Gsk3β was responsible for Ampk inhibition, which accounted for high *de novo* fatty acid synthesis in naïve embryonic stem cells (ESCs). The knockout of glycogen synthase 1 (*Gys1*) in naïve ESCs (GKO), resulting in a drastic glycogen loss, led to a robust Ampk activation and lowered the level of fatty acids. GKO lost the cellular characteristics of naïve ESCs and rapidly transitioned to a primed state, as evidenced by a decrease in pluripotency markers in teratoma from GKO. The characteristics of GKO were restored by the simultaneous knockout of Ampk. These findings suggest that glycogen in naïve ESCs within the blastocyst may act as a signaling molecule for the timely activation of Ampk, thus ultimately contributing to the transition to the epiblast stage.

**Graphical Summary:** 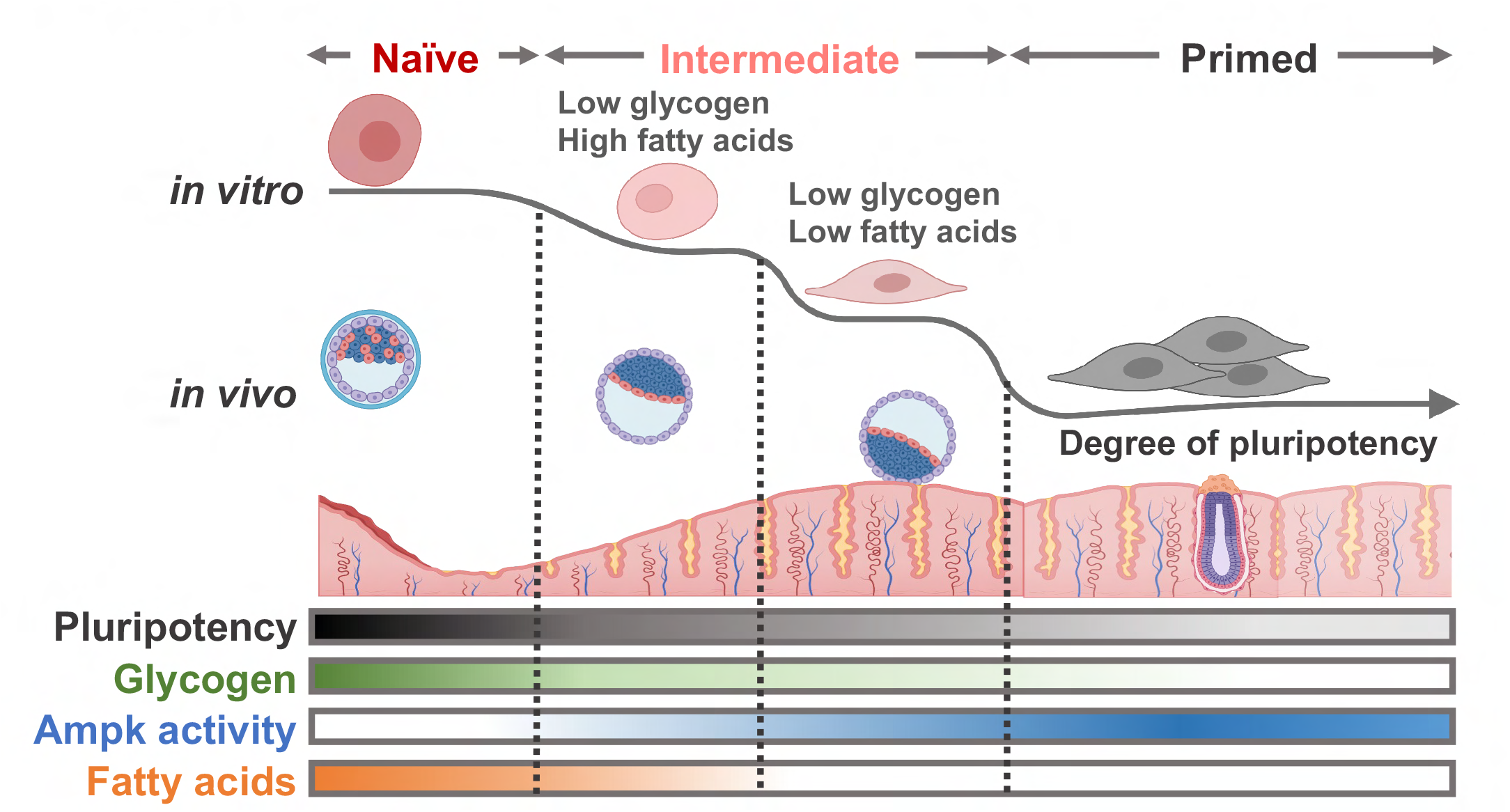

## Introduction

The unique cellular characteristics of naïve and primed embryonic stem cells (ESCs) such as signaling, epigenetics, and metabolism reflect the biology of embryonic and epiblast stem cells of pre- and post-implantation embryos ^1–3^. The requirement of two chemical inhibitors (2i) for Mek1 and Gsk3β pathways for maintaining naïve ESCs epitomizes the constant suppression of Mek1 and Gsk3β pathways by Netrin-1 signaling in the preimplantation embryo ^4^. Additionally, the high dependency of fatty acids on the survival of ESCs ^5, 6^ illustrates the critical roles of fatty acids during early embryogenesis, as demonstrated by the embryo lethality caused by genetic perturbation of key genes involved in fatty acid synthesis such as fatty acid synthase (*Fasn*) ^7^ or acetyl-CoA carboxylase 1 (*Acc1*) ^8^. Thus, the active production of fatty acid up to the blastocyst stage ^9^ suggests that embryos favor fatty acid oxidation (FAO) as a source of energy ^10^.

Particularly, the dynamic change of glucose metabolism during embryo development has also been closely characterized by comparing naïve and primed ESCs (reviewed in ^11, 12^). Although primed ESCs and human induced pluripotent stem cells (iPSCs) predominantly depend on glycolysis ^13^, a clear metabolic shift to bivalent metabolism with capacity for oxidative phosphorylation (OxPHOS) occurs in naïve ESCs ^14, 15^. Such metabolic transition is induced by epigenetic changes from differentially produced metabolites such as *S*-adenosyl methionine ^14^ and α-ketoglutarate ^16^. Interestingly, the metabolic transition from primed to naïve ESCs is initiated by simultaneous inhibition of Mek1 and Gsk3β with 2i ^17–19^ through the upregulation of *Esrrb* ^20^ or *LIN28* ^21^. *Esrrb* expression is a downstream event of Gsk3β inhibition ^22^, and therefore signaling pathways altered by 2i determine the acquisition of naïve pluripotency.

Glycogen dynamics during embryonic development have been characterized for decades ^23, 24^ and its role as a stored energy source has been previously reported ^25^. Particularly, glycogen levels increase with embryonic development from the 2-cell stage to the early blastocyst stage, which is followed by a sharp decrease after the late blastocyst stage ^23, 26^. Although Gsk3β phosphorylates and inhibits glycogen synthase (encoded by *Gys1*), exposure to a Gsk3β inhibitor to consequently activate glycogen synthase only promotes glycogen synthesis in naïve but not primed ESCs ^17^, which is reproduced in human ESCs ^27^. These studies suggest that naïve specific glycogen itself has roles in maintaining naïve pluripotency ^17, 27^.

This study aimed to determine the role of intracellular glycogen, which we had previously found to exclusively occur during naïve pluripotency. Through the establishment of a *Gys1* knockout, we demonstrated that intracellular glycogen would serve as a signaling molecule for the timely activation of Ampk to repress the *de novo* synthesis of fatty acids, whose temporal reduction leads to an exit from naïve pluripotency.

## Material and Methods

### Cell culture

Naïve mouse ESCs were cultured on 0.5% porcine gelatin-coated dish in Naïve mESC culture media -DMEM high glucose (Gibco) supplemented with 15% FBS (Gibco), 1% Glutamax (Gibco), 1% MEM-nonessential amino acids (Gibco), 0.1% Gentamycin (Gibco), 0.1 mM β-mercaptoethanol (Gibco), 1,000 U/ml mouse leukemia inhibitory factor (mLIF) (Millipore, Merck), 1 μM PD0325901 (Peprotech) and 3 μM CHIR99021-at 37 °C and 5% CO_2_ incubating condition. Cells were passaged 1:20 ratio every 3 days using 0.25% Trypsin/EDTA (Wellgene) as a single cell dissociation reagent. OG2+/-GOF6+/-cells were cultured on 0.5% porcine gelatin-coated dish in Naïve mESC culture media with or without 2i (1 μM PD0325901 (Peprotech) and 3 μM CHIR99021). Primed mouse ESCs were cultured on Matrigel (Corning# 354277)-coated dish in EpiSC culture media -DMEM/F12 (Gibco) supplemented with 20% KnockOut Serum Replacement (Gibco), 1% GlutaMAX (Gibco), 1% MEM-nonessential amino acids (Gibco), 0.1% Gentamycin (Gibco), 10 ng/ml murine bFgf (Peprotech), 20 ng/ml murine Activin (Peprotech)-at 37 °C and 5% CO_2_ condition. Primed mESCs were passaged every 3 days using 1 U/ml of Dispase (Gibco) as a colony detachment reagent. Detached colony clumps were transferred 1:15–1:20 ratio on Matrigel (Corning# 354277) coated dishes. Culture media was changed every day for all cell types.

### Establishment and sequence validation of Knock Out cell lines

For establishment of Gys1 KO mESCs, we transfected pRGEN-Cas9-CMV-Puro-RFP (Toolgen-TGEN_OP1) plasmids with sgRNA of Gys1 cloned into pRG2 (Addgene# 104174) plasmid with following sgRNA sequence (5’-GGACACAGCCAATACAGTCA-3’). The cloned gRNA vector (3 μg) and Cas9 plasmid vector (1 μg) were co-transfected into 1x10^6^ mESCs using Lipofectamine 3000 reagent (#L3000-001, Invitrogen). After 24 hours, cells were selected with puromycin (2 μg/mL) for 24 hours followed by washing with culture media. Single-colony picking was performed from KO pool. Targeted sequence of each single clone was validated by Sanger sequencing after gDNA isolation through Wizard® Genomic DNA Purification Kit (#A1120, Promega). For *Prkaa1* knockout, commercial CRISPR/Cas9 Knockout Plasmid (#sc-430618, Santa Cruz) and HDR plasmid (#sc-430618, Santa Cruz) were co-transfected using Lipofectamine 3000 reagent (#L3000-001, Invitrogen). After 24 hours, RFP positive cells were sorted using FACS Aria III cell sorter (BD Biosciences).

### T7E1 assay

KO pool cells were collected and gDNA was extracted using Wizard® Genomic DNA Purification Kit (#A1120, Promega) following the manufacturer’s instruction. T7E1 assay was performed as previously described ^28^.

### Establishment of Knock Down cell lines

For mouse shGys1 plasmid vector generation, we used PB-PGK-Neo empty vector as a backbone vector and the shRNA sequence for mouse Gys1 was infusion cloned into the plasmid vector (shGys1 sequence, forward: 5’CAAGGGTTGTAAGGTGTATTTCTCGAG3’, reverse: 5’ CAAGGGTTGTAAGGTGTATTTCTCGAG3’). To establish *Gys1* stable knock-down cell line, 2 μg of sh*Gys1* Piggy-Bac vector and 1 μg of Transposase vectors was co-transfected into 1x 10^6^ cells of mESCs through Lipofectamine-3000 transfection (#L3000-001, Invitrogen). After 24 hours, cells were treated with 100 μg/mL of G418 (Sigma) for 2 days followed by washing-off G418 on following day.

### Immunoblotting analysis

RIPA buffer (Biosesang) containing 1 μM protease inhibitor and 10 μM sodium orthovanadate was used to extract the whole cell lysate, which was then acquired after incubating on ice for 1 hour and subsequent centrifugation. Quantification of proteins was performed using the Pierce BCA protein assay Kit (#23225, Thermo Fischer Scientific). To prepare the protein sample, 5× SDS-PAGE loading buffer (#SF2088-110-00, Biosesang) was added, and the sample was boiled at 100 °C for 10 minutes. 10μg∼20μg of total protein was loaded and separated on a 10% SDS-PAGE gel. The separated proteins were transferred onto an activated PVDF membrane. The membrane with transferred proteins was blocked with 5% skim milk in TBS-T at RT for 1 hour, followed by washing. The primary antibody (1:500 ∼ 1:1000) in TBS-T was incubated with 1% sodium azide at 4 °C overnight. After washing, the membrane was incubated with the secondary antibody (1:10000) in TBS-T at RT for 1 hour. Chemiluminescence was detected using the Miracle-Star (#16028, iNtRON Biotechnology) kit or West-Queen (#16026, iNtRON Biotechnology) kit.

### RNA isolation and quantitative RT-qPCR analysis

The Easy-Blue™ total RNA isolation kit (iNtRON Biotech) was used to isolate total RNA from cells, following the manufacturer’s instructions. During reverse transcription, 5× PrimeScript™ RT mix (TaKaRa) was used to acquire cDNA. Quantitative real-time PCR was carried out using 2× TB-Green premix (TaKaRa) on a LightCycler-480®II (Roche). The Rn18s gene was used as an internal loading control to normalize the gene expression data. Primer sequence information of each gene for RT-qPCR is listed in Supplementary Table 1.

### CDg4 and Nile Red staining and quantification

For CDg4 and Nile Red staining of *in vitro* cultured cells, cells were dissociated with Accutase (#561527, BD Bio-sciences) and washed with DPBS. 1 million cells were counted followed by fixation with 4% PFA at RT for 5min. After fixation, cells were incubated with CDg4 dye (3μM) or 1 μg/mL of Nile Red in DPBS at 37°C for 1 hour. After washing with DPBS, each fluorescent dye was analyzed by Flow Cytometry. For blastocysts staining, 3.5-4.5dpc mouse embryos were collected fixed with 4% PFA followed by 3μM of CDg4 or 1 μg/mL of Nile Red staining for 1hr at 37°C. 10 μg/mL of Hoechst was counter stained to indicate nucleus of each embryo. The fluorescent and bright field images of the embryo were taken after the staining.

### BODIPY 493/503 staining and quantification

For BODIPY 493/503 staining of *in vitro* cultured cells, cells were dissociated with Accutase (#561527, BD Bio-sciences) and washed with DPBS. 1 million cells were counted and incubated with BODIPY 493/503 (2μM) in DPBS at 37°C for 30 min. After washing with DPBS, BODIPY 493/503 staining was analyzed by Flow Cytometry.

### Flow cytometric analysis

Flow cytometric analysis was used to measure the GFP intensity after BODIPY 493/503 and CDg4 staining, as well as GFP and RFP intensity of OG2GOF6 cell line. Cells were dissociated with Accutase (#561527, BD Bio-sciences) and washed with DPBS. Cells were analyzed through FACS Calibur (BD Biosciences) or FACS Celesta (BD Biosciences). GFP and CDg4 intensity was determined by measuring FL-1 channel (FACS Calibur) or FITC (FACS Celesta). For RFP and Nile Red intensity, FL-3 channel (FACS Calibur) was measured. FlowJo software was used for analyzing the flow cytometric data.

### NMR analysis with ^13^C-glucose tracer

Cells were cultured with 15 mM ^13^C-glucose (U-^13^C_6_, 99%, Cambridge Isotope Laboratories) for 24 hours. After harvesting, cell pellets underwent standard two-phase extraction, and the chloroform phase underwent speedvac. The dried samples were dissolved in chloroform-d_6_ (Sigma-Aldrich) and ^1^H-^13^C Heteronuclear Single Quantum Coherence (HSQC) NMR was taken, using 800 MHz Bruker Avance III HD spectrometer equipped with a 5 mm CPTCI CryoProbe (Bruker BioSpin, Germany). The time domain parameter for carbon was 512, the spectral width for carbon was 80, and the number of scans was 3.

### Cell imaging

Brightfield images of live cells were captured by Olympus CKX41. GFP and RFP images of live cells were captured by JuLi Stage (NanoEntek), followed by the merging process of each channel by JuLi-Edit software.

### Glycogen quantification assay

Glycogen assay kit (BioVision) was used to measure the relative intracellular glycogen amount. For the sample preparation, cells were dissociated with Accutase (#561527, BD Bio-sciences) and washed with DPBS. 3 million cells were homogenized with cold distilled water on ice for 1 hour. Supernatant was collected after spin down (13000 rpm, 20 min). BCA assay was performed with the lysate. Then the lysate was boiled at 100 °C for 10 min. 30∼50 μg of protein was loaded to each well of 96 well plate and glycogen assay was performed under the manufacturer’s instruction. The optical density of glucose (which was hydrolyzed from glycogen) was measured at 570nm by Epoch microplate spectrophotometer (Biotek). Final glycogen concentration was normalized into initial protein input of each sample.

### Teratoma formation assay

To conduct the teratoma formation assay, ten 3-week-old male BALB/C nude mice were housed in the mouse facility for one week to acclimate before undergoing cell injection. One million cells of each line, Cont and GKO mESCs, were prepared in 100 µL of mESC media for the assay. The cells were then injected into the mice either intratesticular (IT) or subcutaneously (SC) after anesthesia. After three weeks from the injection, the mice were euthanized and the size of each teratoma tissue was measured. Each teratoma volume was measured as following formula: V (cm^3^) = a (cm) × b (cm) × c (cm) × π / 6 (a = length, b = width, and c = depth). The tissues were subsequently isolated for RNA and subjected to RT-qPCR analysis. Animal experiments were conducted under the permission of Seoul National University Institutional Animal Care and Use Committee (Permission number: SNU-211001-1-2)

### Bulk RNA-seq library preparation

Total RNA was isolated from J1, GKO and PJ1 cellusing Easy-BLUETM RNA isolation kit (iNtRON Biotechnology, #17061). One 1 µg of total RNA was processed for preparing mRNA sequencing library using MGIEasy RNA Directional Library Prep Kit (MGI) according to manufacturer’s instruction. The first step entails utilizing poly-T oligo-attached magnetic beads to isolate the mRNA molecules that contain poly-A. Following purification, divalent cations and a high temperature are used to break the mRNA into small pieces. Utilizing reverse transcriptase and random primers, the cleaved RNA fragments are converted into first strand cDNA. After achieving strand specificity in the RT directional buffer, second strand cDNA synthesis takes place. The ’A’ base is then added to these cDNA fragments, followed by the ligation of the adapter. The final cDNA library is made by purifying and enriching the results with PCR. The QauntiFluor ONE dsDNA System (Promega) is used to quantify the double stranded library. The library is circularized at 37 °C for 30 min, and then digested at 37 °C for 30 min, followed by cleanup of circularization product. The library is treated with the DNB enzyme at 30 °C for 25 min to create DNA nanoballs (DNB). Finally, Library was quantified by QauntiFluor ssDNA System (Promega). On the MGIseq system (MGI), the prepared DNB was sequenced using 100 bp paired-end reads.

### Bulk RNA-seq processing and analysis

Low-quality bases and adapter sequences bases were trimmed using TrimGalore (https://www.bioinformatics.babraham.ac.uk/). The trimmed reads were aligned to the mouse genome assembly GRCm39 using STAR (v2.7.3a). The expression value per gene was estimated as a read count or transcripts per million (TPM) values calculated using RSEM (v1.3.3) based on the mouse gene annotation GRCm39.104. The specific gene set enrichment variation was assessed by gene set variation analysis (GSVA) using the R package.

### Re-analysis of published Phosphoproteome data

The processed data from Ana Martinez-Val et al’s study was analyzed using a two-sided limma approach (version 3.54.2) to determine differential phosphorylation sites at each time point compared to the initial Serum/LIF conditions. The statistical threshold was adjusted using the Benjamini-Hochberg correction to control the false discovery rate (FDR) at 5% for each condition. The PHONEMsS tool was used to model signaling networks using perturbed phosphorylation sites. Further kinase enrichment analysis was performed using Fisher’s exact test ^29^.

### Statistical Analysis

The mean values of the quantitative data were presented with their corresponding standard deviation (SD). To determine the statistical significance of each response variable, unpaired two-tailed t-tests were performed. Where necessary, pre-specified comparisons between groups were conducted using Tukey’s post hoc test in PRISM software. *p*-values less than 0.05 were considered statistically significant (* < 0.05, ** <0.01, ***<0.001, ****<0.0001 and n.s. for not significant).

## Results

### Reduced Ampk-dependent phosphoproteome during the transition to the naïve state with 2i

The induction of naïve pluripotency to mimic the pre-implantation embryos can be achieved through simultaneous inhibition of Mek1 and Gsk3β with chemical inhibitors (hereinafter referred to as 2i) along with leukemia inhibitory factor (LIF). The timely alteration of the transcriptome during this transition provides a snapshot of the changes in gene expression patterns in response to these agents ^30^. However, the kinase signaling pathways that initiate such gene response upon LIF/2i treatment remain largely unexplored. To this end, we took advantage of a recently published phosphoproteome dataset collected at multiple stages of naïve pluripotency induction ^31^, as shown in Figure 1A. Through a meta-analysis of the phosphoproteome datasets, we identified four clusters with distinct phosphoproteome signatures (SL: Cont, 0.5hr, 1hr/2hr, and 6hr) (Fig. 1B). The differentially phosphorylated proteins (DPs) became manifested from 0.5hr after 2i (Fig. 1C) to 6 hours after 2i treatment (Extended data Fig.1). The signal network models based on kinase/phosphatase-substrate interactions ^32^ revealed that reduced phosphorylation of downstream substrates of Mapk1 (i.e., Erk2) and Gsk3β was manifested at 0.5hr after concurrent inhibition of Mek1 (with iMek1) to Erk1/2 and Gsk3β (with iGsk3β) (Fig. 1D). The subsequent kinase enrichment analysis from reduced DPs uncovered the AMPK pathway along with MAPK1 and ERK1 (Fig. 1E). The temporal profile of DPs also validated the 2i treatment in parallel with the inhibition of MAPK1 and GSK3 signaling (Fig. 1F), which eventually inhibited AMPK signaling. Particularly, reduced DPs are among the typical substrates for AMPK (Fig. 1G).

**Figure 1.**
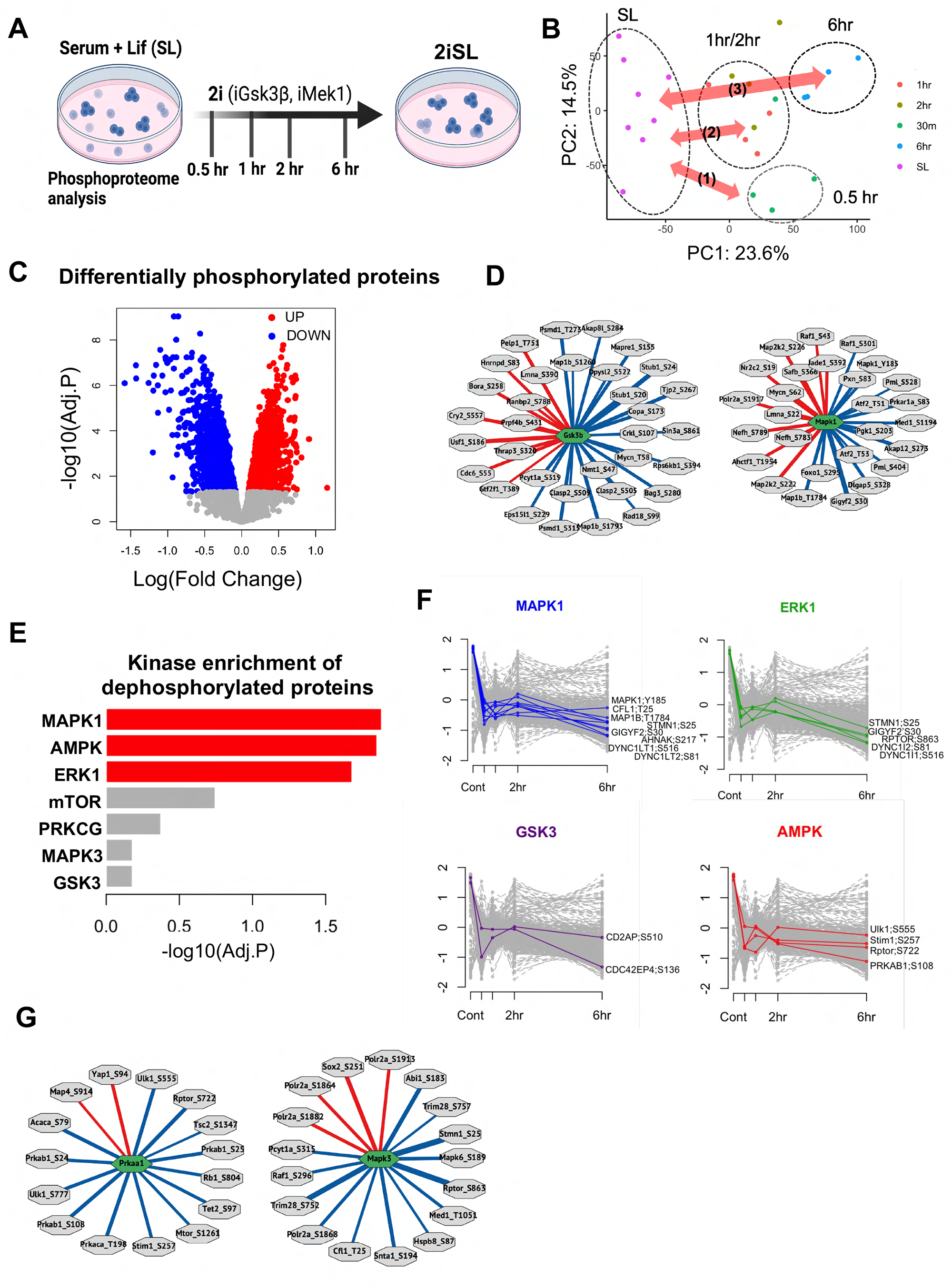
(A) Graphical image of experiment from Mariniez-Val et al. (B) Principal Component Analysis of the phosphoproteome from (A). (C) Volcano plot of differentially phosphorylated proteins after 30min of treatment compared to SL. (D) PHONEMeS-ud solution model of signaling of GSK3B(Gsk3b) and MAPK1(Mapk1) after 30min of 2i treatment. Green hexagons correspond to target proteins and gray octagons correspond to substrates of target proteins. A red bar represented an upregulated of phosphorylation and a blue bar represented a downregulated of phosphorylation. (E) Kinase analysis of down-regulated phosphoproteins over time. Selected kinases are highlighted. p-value<0.05, Fisher’s exact test. (F) Clustering of phosphosites based on their temporal dynamics. Four clusters enriched for known substrates of MAPK (blue), ERK1 (green), GSK3 (purple), and AMPK (purple) are shown. (G) PHONEMeS-ud solution model of signaling of AMPK(Prkaa1) and MAPK3(Mapk3) after 30min of 2i treatment.

### A high level of fatty acid in naïve ESCs is concurrent with Ampk inhibition

Based on the drastic decrease of the Ampk-dependent phosphoproteome by 2i treatment (Fig. 1), we next monitored Ampk activity between the isogenic pair of naïve and primed ESCs, two types of cells that have been previously described ^2, 17, 18, 33^ (Fig. 2A). As expected, the active phosphorylation of Ampk was manifested in primed ESCs in parallel with Erk1/2 phosphorylation (due to supplement of Fgf2) and loss of Stat3 phosphorylation (due to lack of LIF) compared to naïve ESCs (Fig. 2B). The active Ampk in the primed ESCs was validated by inhibitory phosphorylation of acetyl CoA carboxylase (Acc), a typical downstream target protein of Ampk ^34^ (Fig. 2C). It is also worth noting that Acc activity is critical not only for *de novo* fatty acid synthesis but also fatty acid β-oxidation (FAO), which accounts for the significant reduction of total fatty acid determined by Nile Red staining in primed ESCs compared to naïve ESCs (Fig. 2D). To identify the mechanisms that led to the reduction of fatty acids in primed ESCs, we next examined the *de novo* fatty acid synthesis using 13C glucose (Fig. 2E). The levels of fatty acids that were newly synthesized from glucose, as determined the split peaks of the omega methyl signal on the NMR spectrum ^35^, were much higher in naïve ESCs (Fig 2F). The deprivation of glucose significantly reduced the intensity of Nile Red staining in naïve ESCs, suggesting that active *de novo* fatty acid synthesis is responsible for high fatty acid levels (Fig. 2G). The higher reactivity of the inner cell mass of E3.5 mouse blastocysts to Nile Red also further supports the exclusive occurrence of fatty acids in naïve ESCs (Fig. 2H). Consistently, inhibition of fatty acid synthesis with the chemical inhibitor of Acc induced drastic cell death in naïve ESCs (Fig. 2I).

**Figure 2.**
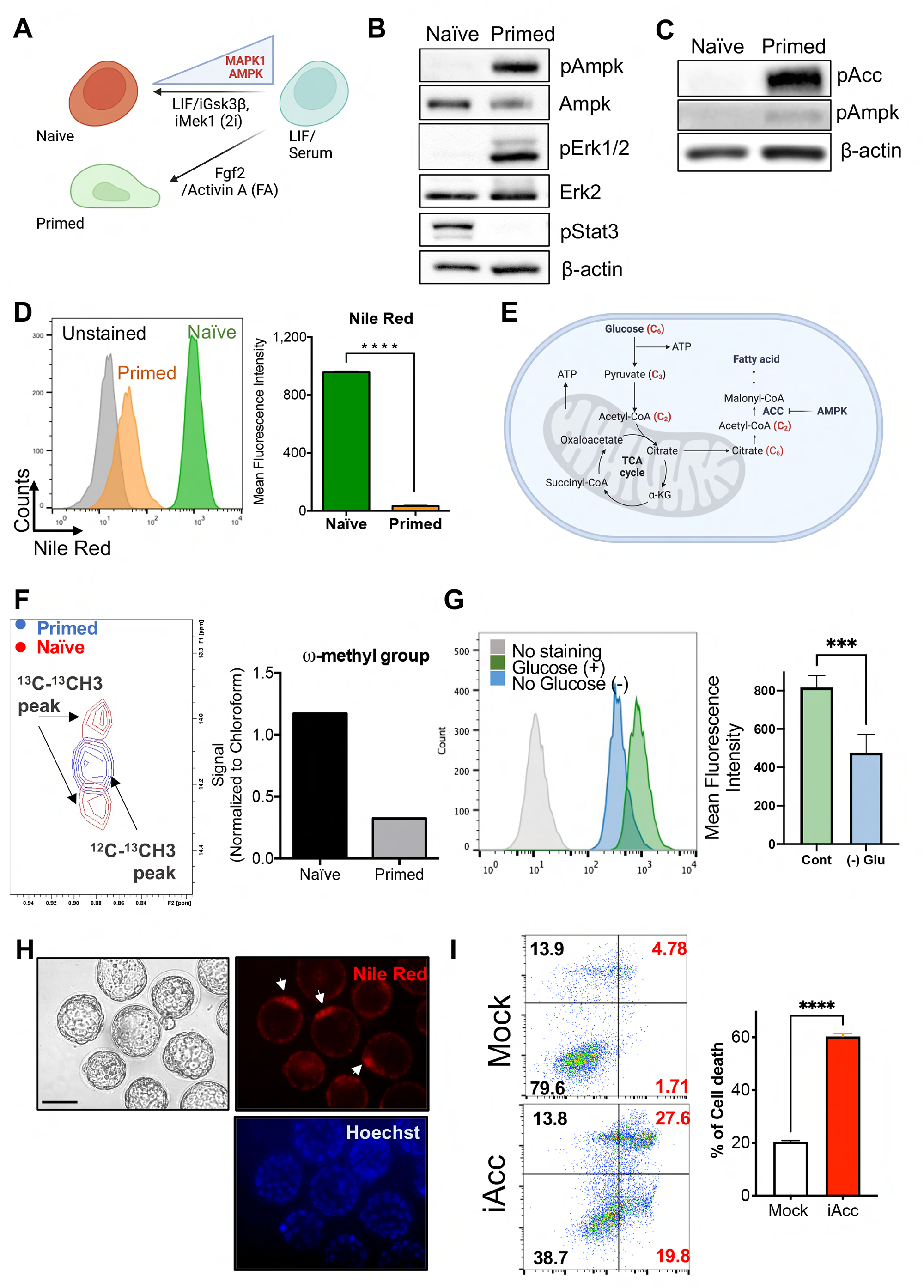
(A) Graphical abstract of naïve conversion (LIF/iGsk3β/iMek1 [2i]) and primed conversion (Fgf2/ActivinA [FA]) from intermediate status (LIF/Serum). (B) Immunoblotting analysis for indicative proteins (pAmpkα, Ampk, pErk1/2, Erk2, pStat3 in J1 (naïve) and PJ1 (primed) mESCs), *β-actin was used as a loading control. (C)* Immunoblotting analysis for indicative proteins (pACC, pAmpkα in J1 (naïve) and PJ1 (primed) mESCs), β-actin was used as a loading control. (D) Flow cytometry of Nile Red staining of J1 (naïve) and PJ1 (primed) mESCs (left panel), quantification of mean fluorescence intensity from the flow cytometry (right panel). (E) Graphical abstract of overall glucose metabolism containing glycolysis, TCA cycle and de novo fatty acid synthesis. (F) (Left) ^1^H-^13^C Heteronuclear Single Quantum Coherence (HSQC) NMR spectra for Primed (Blue) and Naïve (Red) cells in the region of ω-methyl group of fatty acids. The split peaks indicate the last two carbons of the fatty acid tail are all ^13^C-labeled. (Right) The signal intensity of the split peaks in Primed and Naïve cells in the left, normalized by the signal intensity of the solvent chloroform of each sample. (G) Flow cytometry of Nile Red staining of J1 with [Cont] or without [(-)Glu] glucose. Glucose was depleted for 24 hrs before the analysis and quantification of mean fluorescence intensity from the flow cytometry (right panel). (H) Mouse embryos (3.5-4.5dpc) were collected fixed with 4% PFA followed by 1 μg/mL of Nile Red suspended for 1hr at 37°C. Hoechst was used for nucleus counterstaining of embryos. Scale bar, 100 μm. (I) Cell viability after iAcc treatment on mESCs was measured through flow cytometric analysis of Annexin-V/7-AAD staining.

### High intracellular glycogen represses Ampk activation in naïve ESCs

The requirement of both iMek1 and iGsk3β to maintain naïve pluripotency ^36^ was previously highlighted, as deprivation of either iMek1 or iGsk3β disrupts the typical dome-shape morphology of naïve ESC colonies ^2^. As Ampk is a downstream substrate of both Mek1/Erk1 and 2 ^37^ and Gsk3β ^38^, we next sought to identify the specific kinase responsible for activating Ampk in ESCs. To this end, naïve ESCs were subjected to depletion of either iMek1 or iGsk3β, and Ampk activity was monitored. While Ampk activation was observed upon withdrawal of iGsk3β in naïve ESCs (Fig. 3A), no such effect was seen in primed ESCs (Fig. 3B). In contrast, iMek1 depletion had a marginal effect on Ampk (data not shown). The inhibition of Ampk by iGsk3β in naïve ESCs was intriguing, as it has been previously reported that Gsk3β directly phosphorylates Ampk, leading to its inhibition in somatic cell models ^38^. This discrepancy suggested the involvement of other factors in the observed inhibition of Ampk with iGsk3β specifically in naïve ESCs (Fig. 3B).

**Figure 3.**
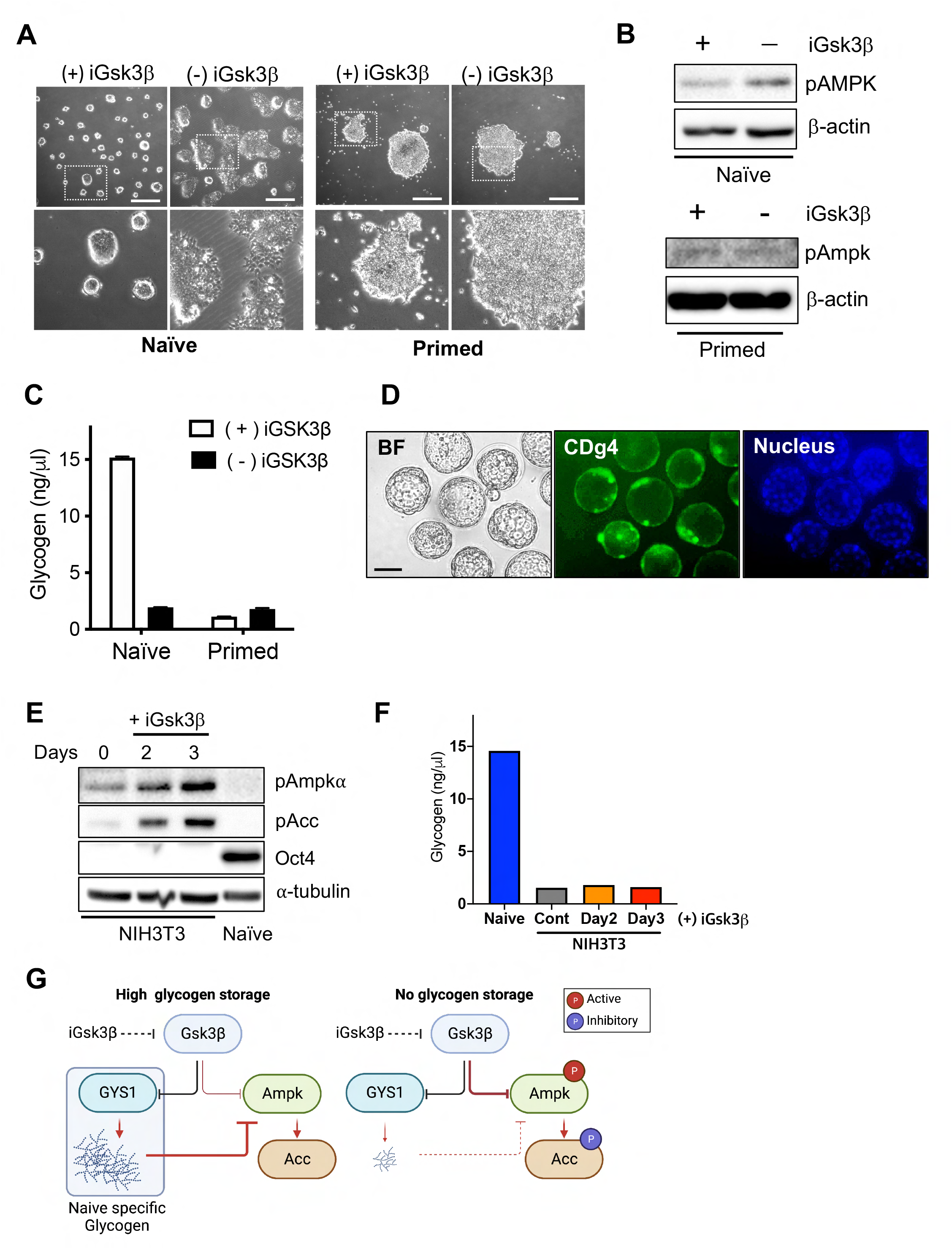
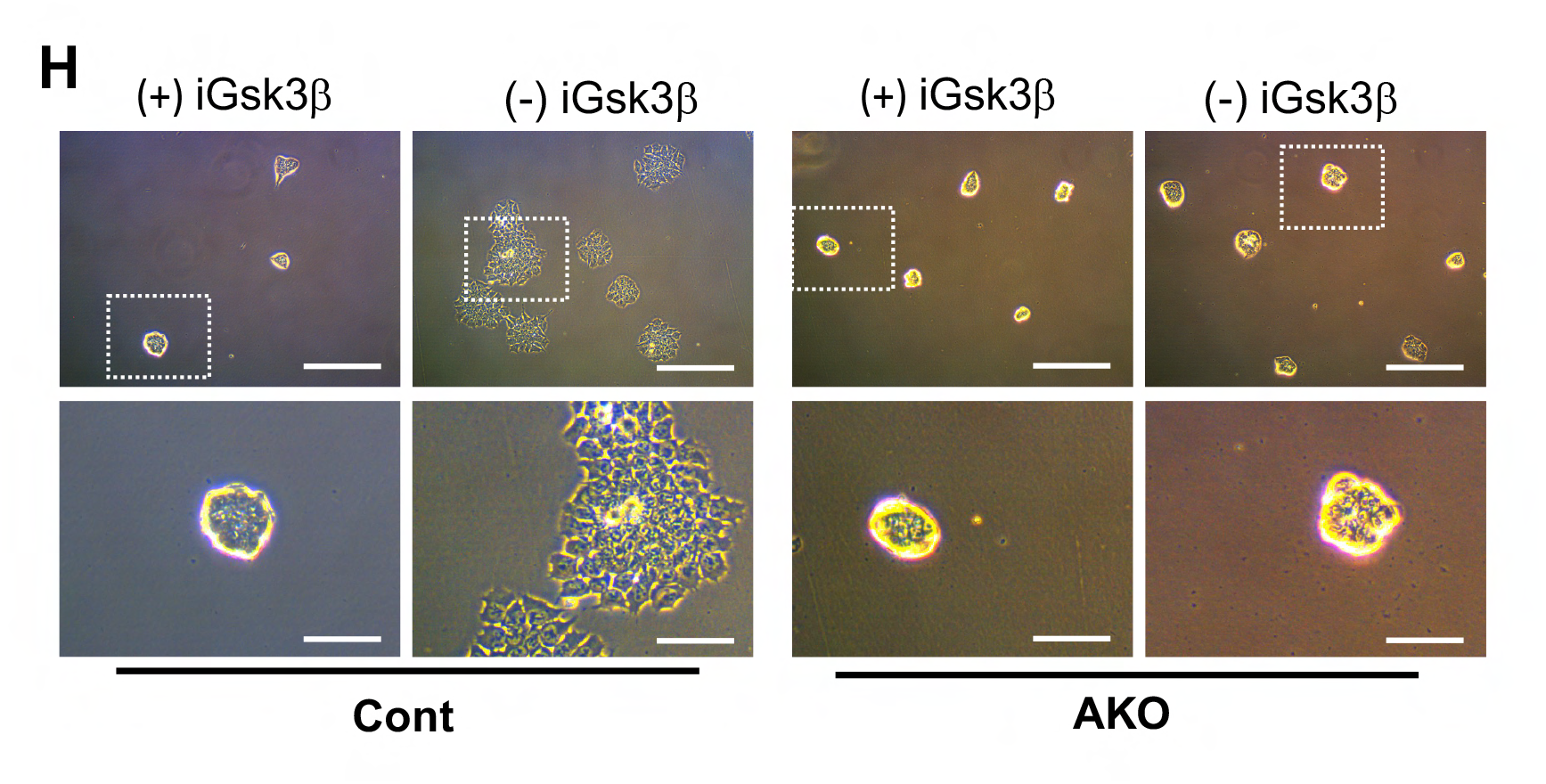
(A) Representative brightfield images of J1 (naïve) and PJ1 (primed) mESCs with or without iGsk3β treatment (scale bars *=* 500 μm). (B) Immunoblotting analysis for indicative proteins (pAmpkα in J1 (naïve) and PJ1 (primed) mESCs with or without iGsk3β treatment). *(C)* Representative brightfield images of J1 Cont and J1 AKO mESCs with or without iGsk3β treatment (scale bars = 500 *μm)*. (D) Intracellular glycogen level of J1 (naïve) and PJ1 (primed) mESCs with or without iGsk3β treatment. (E) 3.5-4.5dpc of mouse blastocysts were collected and fixed with 4% PFA. 3 μM of CDg4 was treated for 1hr for embryo staining. Hoechst for nucleus counterstaining. Scale bar, 100 μm. (F). Immunoblotting analysis for indicative proteins (pAmpkα, pAcc, Oct4 in J1 (naïve) and NIH3T3 under indicative conditions), α-tubulin was used for the loading control. (G) Intracellular glycogen level of J1 (naïve) and NIH3T3 under indicative conditions. (H) Graphical abstract of intracellular signaling mechanism for Ampk and Acc activity under high glycogen storage condition (left panel) or no glycogen storage condition (right panel). Cont: Control, AKO: *Prkaa1* KO.

On the other hand, previous studies have reported that intracellular glycogen can inhibit Ampk activity by directly interacting with the glycogen binding domain (CBD) of the Ampkβ subunit ^39^. Notably, unlike primed ESCs, naïve ESCs have been shown to produce glycogen upon Gsk3β inhibition ^17^. Accordingly, depletion of the Gsk3β inhibitor markedly reduced stored glycogen in naïve ESCs but not in primed ESCs (Fig. 3C). This suggests that the production of glycogen in naïve ESCs upon Gsk3β inhibition may be a contributing factor to the observed inhibition of Ampk activity in these cells (Fig. 3B). Naïve-specific glycogen production by iGsk3β (Fig. 3C) was highlighted by clear staining of CDg4, a fluorescent probe for glycogen ^17, 40, 41^ in the ICM of a mouse blastocyst (Fig. 3D), which was consistent with previous observations ^23^. In sharp contrast to mESCs (Fig. 3E), the activation of Ampk by Gsk3β inhibition (with iGsk3β) occurred in NIH3T3 cells (i.e., normal somatic cells) where glycogen was obviously lacking (Fig. 3F). Thus, we concluded that naïve specific glycogen production by Gsk3β inhibition could explain the contradictory repression of Ampk activity in naïve ESCs (Fig. 3G). Based on the hypothesis that Ampk activation by removal of iGsk3β contributes to the loss of pluripotency, further experiments were conducted using Ampk knockout (AKO) naïve ESCs (Extended data Fig. 2A). Interestingly, the AKO cells retained the typical dome-shaped morphology even after depletion of iGsk3β, whereas the control group largely lost this characteristic dome shape (Fig. 3H). This suggests that Ampk activation may play a role in the loss of naïve pluripotency induced by iGsk3β depletion, and its absence through knockout may prevent the morphological changes associated with loss of naïve pluripotency in response to iGsk3β depletion (Fig. 3H). Consistently, typical naïve marker genes were highly expressed in AKO compared to the control (Extended data Fig. 2B).

### Role of glycogen in the suppression of Ampk activation and maintaining fatty acid levels

We next sought to determine the role of glycogen through Ampk in naïve ESCs. To this end, we first established a line of naïve ESCs lacking glycogen by introducing an indel (insertion and deletion) at exon 9 of glycogen synthase 1 (Gys1) (Fig. 4A). After conducting the T7E1 assay to confirm the knockout of Gys1 in naïve ESCs (Fig. 4B), multiple single clones of Gys1 KO naïve ESCs (hereinafter referred to as GKO) were established. As expected, intracellular glycogen was completely depleted in the GKO (Fig. 4C) and the cells that reacted with CDg4 were markedly reduced (Fig. 4D). The temporary resistance to glucose deprivation in naïve ESCs, which was accompanied by a loss of stored glycogen ^17^, was markedly reduced in GKO (Extended data Fig. 3A), which would be strongly associated to a lack of stored glycogen in GKO (Fig. 4C). In parallel with glycogen depletion, Ampk activation and consequent Acc inhibitory phosphorylation were manifested in GKO (Fig. 4E), suggesting that the intracellular glycogen in naïve ESCs is responsible for the constant inhibition of Ampk. Consistent with the high inhibitory phosphorylation of Acc to impede *de novo* fatty acid synthesis, the fatty acid level was markedly reduced in GKO, as determined by BODIPY staining (Fig. 4F). Therefore, transient glucose withdrawal markedly reduced the level of fatty acids in GKO (Fig. 4G). Additional assays were conducted to rule out the possibility of an unexpected off-target effect of KO, and similar results were obtained through the depletion of Gys1 by stable knockdown (shGys1) in naïve ESCs (Extended data Fig. 3B). The reduced glycogen (Extended data Fig. 3C) and Ampk activation and the accompanying Acc inhibition (Extended data Fig. 3D) coincided with a lower fatty acid level (Extended data Fig. 3E). Ampk activation was manifested in Gys1 knockdown (KD) cells in the presence of iGsk3β, in contrast to the Ampk inhibition observed in control ESCs (Extended data Fig. 3F). These results clearly demonstrated that the intracellular glycogen produced by constant exposure to iGsk3β inhibited Ampk activation and contributed to the maintenance of intracellular fatty acid levels in naïve ESCs.

**Figure 4.**
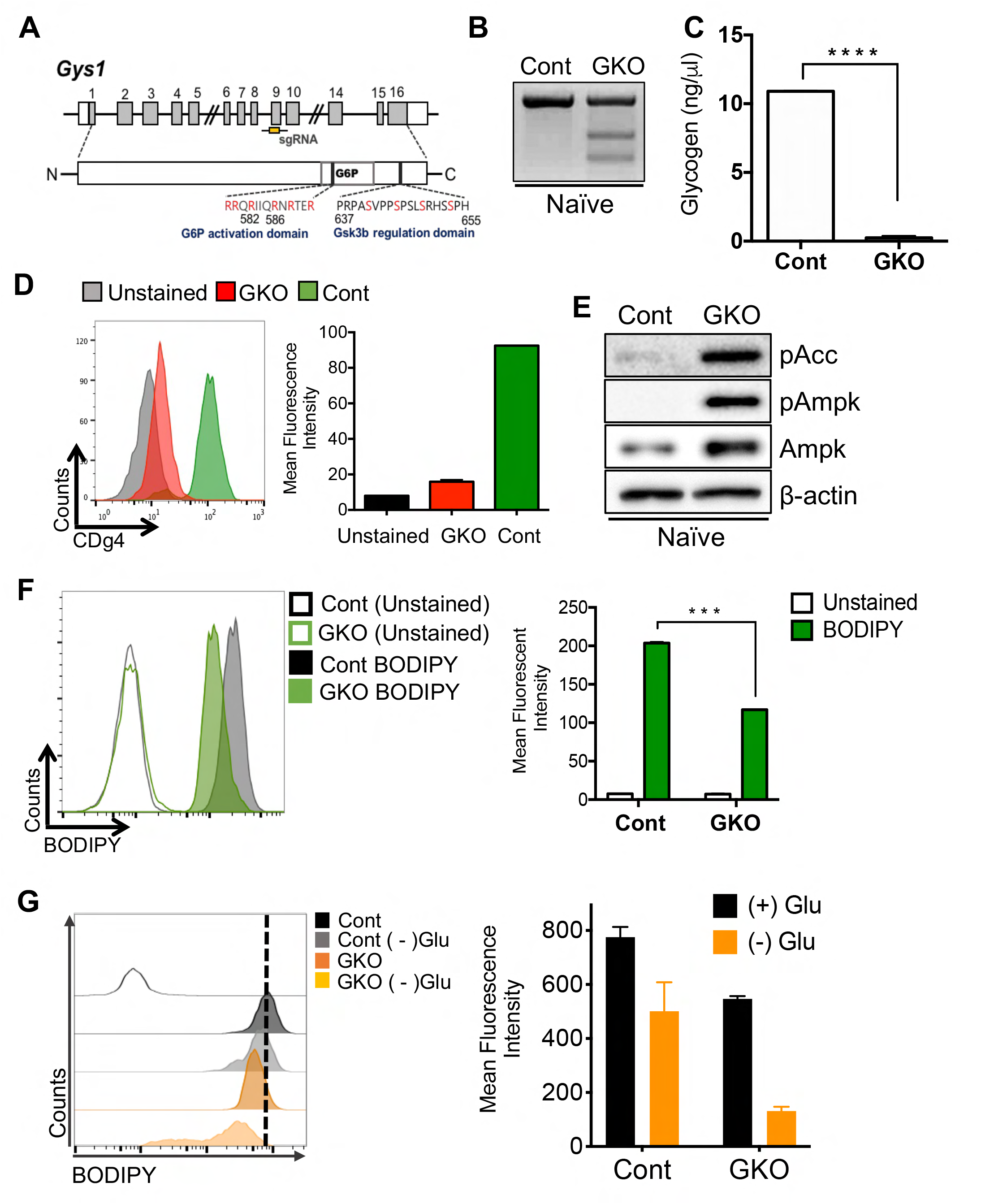
(A) Scheme for *Gys1* KO targeting Exon 9. (B) T7E1 assay for WT and GKO in OG2^+/-^GOF6^+/-^ cell line. (C) Intracellular glycogen level of J1 Cont and GKO. (D) Flow Cytometry of CDg4staining in J1 Cont and GKO (left panel), quantification of the mean fluorescence intensity of the flow cytometry (right panel). (E) Immunoblotting analysis for indicative proteins (pAcc, pAmpkα, Ampk in J1 Cont and GKO), β-actin is used for the loading control. (F) Flow cytometry of BODIPY *493/503* staining in J1 Cont and GKO (left panel), quantification of the mean fluorescence intensity of the flow cytometry (right panel). (G) Flow cytometry of BODIPY *493/503* staining in J1 Cont and GKO with or without glucose (left panel), quantification of the mean fluorescence intensity of the flow cytometry (right panel). Cont: Control, GKO: *Gys1* KO.

### Loss of glycogen primes exit of naïve pluripotency

The distinct cellular difference between the control and GKO cells was the colony morphology. Specifically, the GKO lost the typical dome-shape colony morphology of naïve ESCs (Fig. 5A). The expression levels of typical naïve markers such as *Nanog*, *Rex1*, and *Esrrb* were all reduced in GKO compared to control naïve ESCs even under LIF/2i conditions (Fig. 5B). The transcriptome of the GKO cells exhibited markedly different gene clusters from those of the parent naïve ESCs (Fig. 5C). The gene set of naïve and primed signatures were enriched in Naïve and Primed samples respectively, with GKO cells showing moderate enrichment to all signatures (Fig. 5D). Consistent with these findings, gene sets associated with specific transcription factors downstream of naïve were progressively enriched from naïve, GKO to primed samples (Fig. 5E). It is also worth noting that the dome-shape colony morphology of control ESCs became lost at day 3 after glucose deprivation when the intracellular glycogen was depleted (Fig. 5F), which appeared to be similar to that of GKO prior to glucose deprivation. Prior to cell death induction of GKO after glucose deprivation, a marked induction of primed marker genes such as *Eomes* and *Fgf5* was evident in GKO (Fig. 5G). To further highlight the effect of *Gys1* in the maintenance of naïve pluripotency, ESCs were treated with two distinct fluorescence reporters under the control of the distal (DE) and proximal enhancer (PE) of *Pou5f1*, indicating naïve (expressing green fluorescence protein: GFP) and primed states (expressing red fluorescence protein: RFP), respectively (hereinafter referred to as dual-reporter ESCs) ^42^ (Fig. 5H). Given that an intermediate state of pluripotency can be identified by the occurrence of both GFP and RFP fluorescence, the naïve or primed transition with LIF/2i or Fgf2/Activin A was readily monitored by each fluorescent probe as described previously ^2^. Similar to the KO approach shown in Figure 4A, another GKO line was established in the dual-reporter ESCs (Extended data Fig. 4A). When control and GKO maintained at an intermediate state with LIF only were subjected to LIF/2i, GKO still showed both GFP and RFP unlike the control ESCs, which were promptly converted to a GFP+ population (Figs. 5I and J). This demonstrated that naïve conversion by 2i treatment was impeded when glycogen synthesis was blocked. In other words, glycogen synthesis is necessary for naïve conversion. Next, we performed a comparison of teratomas developed from control and GKO ESCs. Remarkably, the teratomas generated from GKO ESCs were significantly larger than those derived from control ESCs (Extended data Fig. 4B). Additionally, as determined by the level of pluripotency marker genes, the population of undifferentiated cells was visibly reduced in GKO teratomas (Fig. 5K), with less intra-group variation compared to control teratomas (Extended data Fig. 4C). These findings suggest that the differentiation process in GKO ESCs, which lack glycogen, may be more synchronized compared to control ESCs with varying amounts of glycogen.

**Figure 5.**
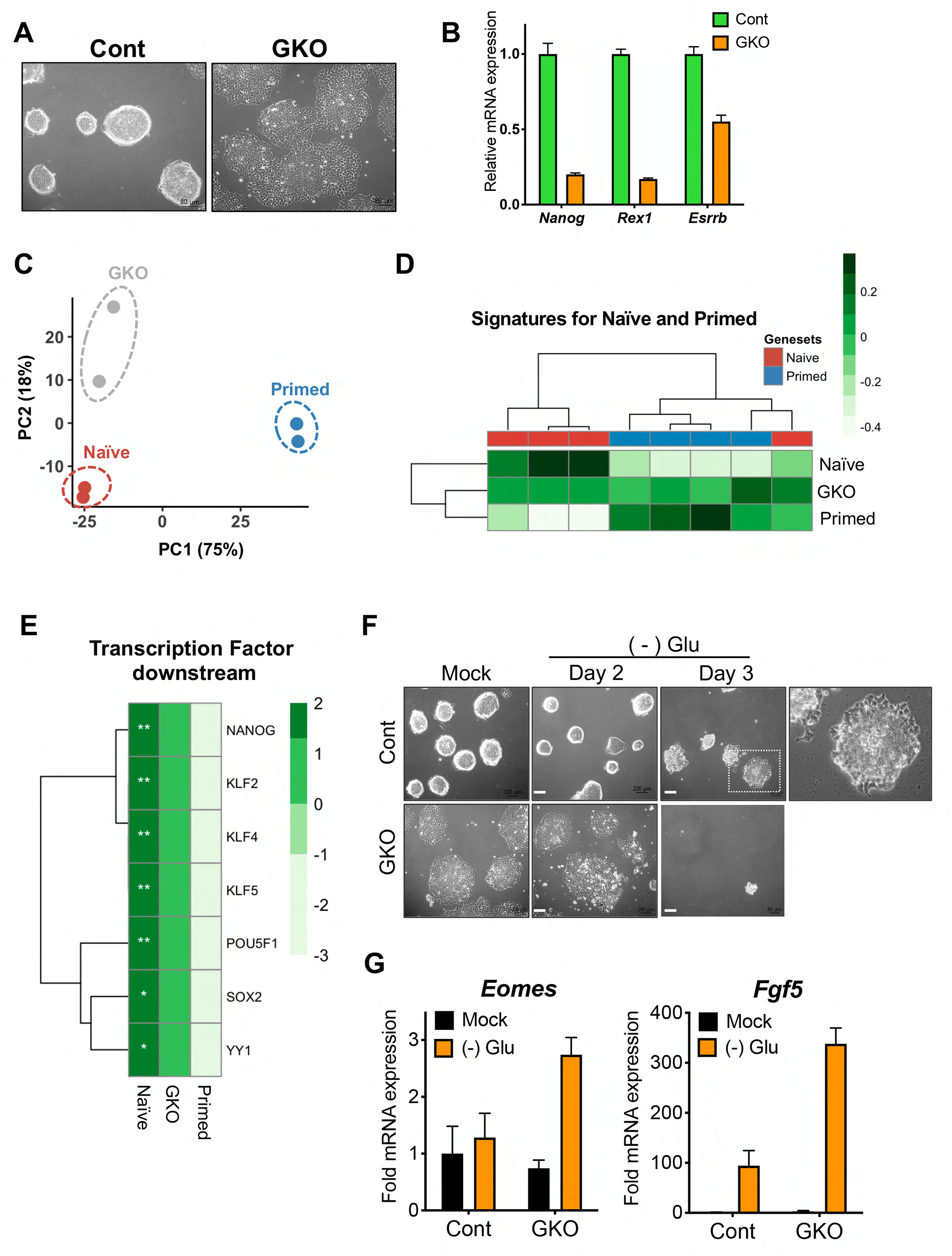
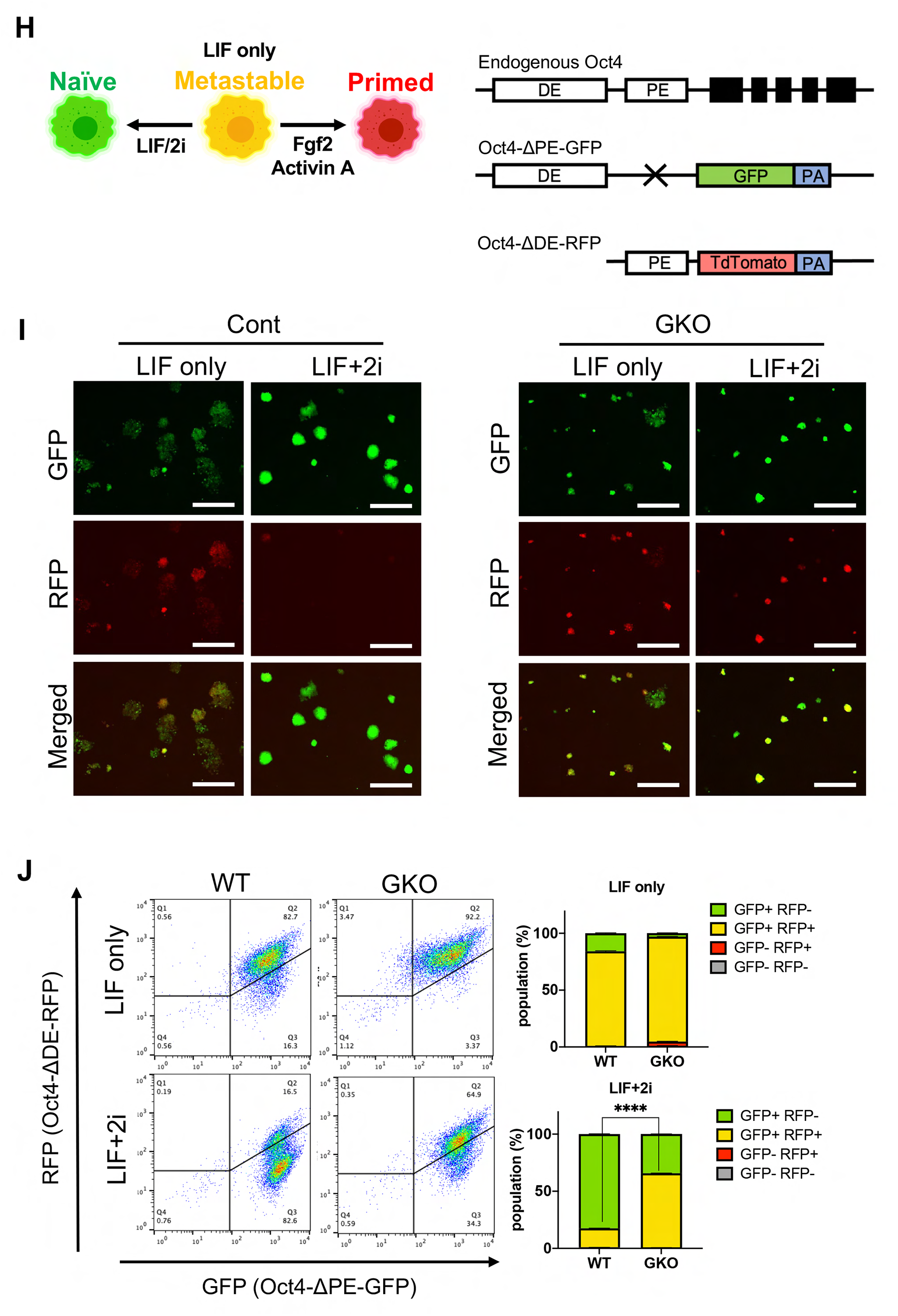
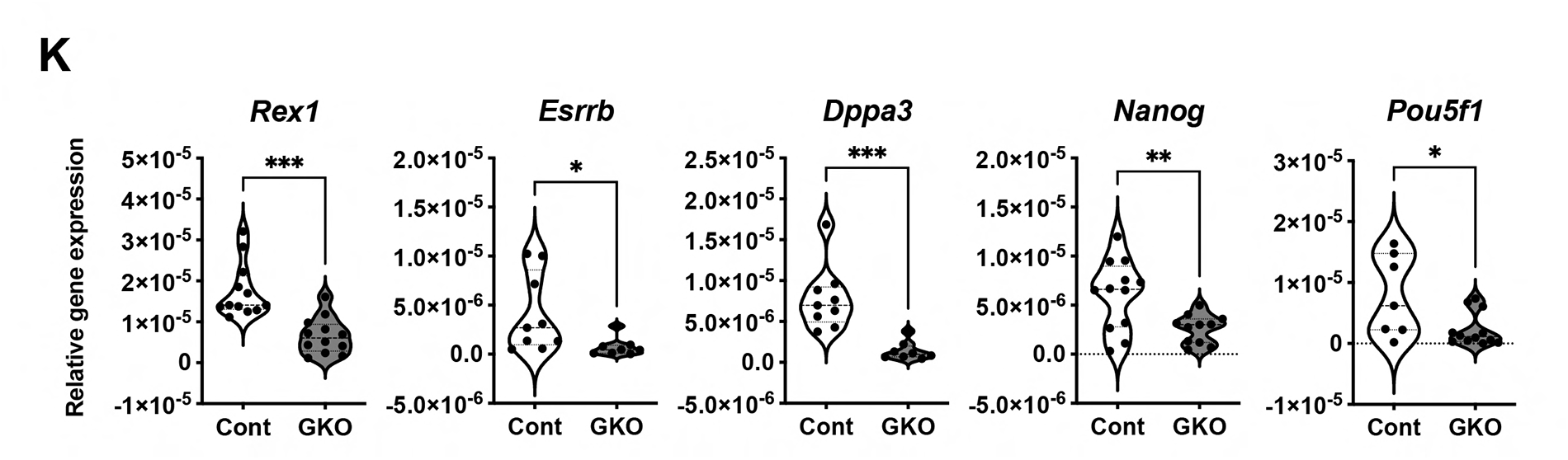
(A) Representative brightfield image of Cont and GKO. (B) Relative mRNA expressions for *Nanog*, *Rex1*, *Esrrb* in Cont and GKO. (C), Principal Component Analysis of the transcriptome of Naïve, GKO and Primed samples. (D) Gene set variation analysis (GSVA) of Naïve and Primed signatures in Naïve, GKO, Primed samples. (E) Gene set variation analysis (GSVA) of Transcription Factor downstream genesets in Naïve, GKO, Primed samples. (F) Representative brightfield image of Cont and GKO with (left) or without (right) Glucose. (G) Fold mRNA expressions of Eomes and Fgf5 in Cont and GKO with [mock] or without [(-) Glu] Glucose. (H) Graphical image of fluorescence activity of OG2^+/-^GOF6^+/-^ cell line in naïve, metastable, primed status (left panel), Graphical scheme of endogenous Oct4 allele, Oct4-dPE-GFP (OG2) allele, and Oct4-dDE-RFP (GOF6) allele (right panel). (I) Representative fluorescence images of Cont and GKO (scale bars = 500μm) under LIF only or LIF+2i conditions. (J) Flow cytometry of GFP and RFP in WT and GKO under LIF only or LIF+2i conditions (left panel), quantifications of the populations from the flow cytometry (right panel). (K) Violin plots of relative mRNA expressions of *Rex1*, *Esrrb*, *Dppa3*, *Nanog* and *Pou5f1* in Teratoma generated from Cont and GKO. (****, p < 0.0001, ***, p < 0.001, **, p<0.01, *, p < 0.05, n = 3), Glu: Glucose. Cont: Control, GKO: *Gys1* KO.

### Ampk is responsible for priming the exit of naïve pluripotency

Our observations that naïve-specific glycogen synthesis by Gsk3β inhibition (Fig. 3) repressed Ampk activation (Fig. 4) and maintained naïve pluripotency (Fig. 5) lead us to hypothesize that Ampk activation, which lowers fatty acid levels, would trigger the exit of naïve pluripotency when glycogen is depleted (Fig. 6A). To demonstrate this hypothesis, Ampk was knocked out through CRISPR/Cas9 gene editing to establish Ampkα KO (AKO) and dual KO of *Gys1* and Ampkα (DKO) (Fig. 6B), as previously described ^43^. The KO ESCs were readily isolated due to the simultaneous integration of RFP reporter through homologous directed repair (Extended data Fig. 5A). Importantly, we actively avoided the use of chemical inhibitors for Ampk perturbation to eliminate the possibility of unexpected side effects by prolonged exposure to these inhibitors, as previously described ^44^. Due to the lack of Ampk in AKO and DKO, inhibitory phosphorylation of Acc in GKO was markedly reduced in DKO (Fig. 6C), which was consistent with the recovery of reduced fatty acids of GKO by loss of Ampk (Fig. 6D) even when glycogen was lacking (Fig. 6E). The clear increase of overall fatty acid levels by AKO demonstrates that Ampk activity determines the level of fatty acids (Fig. 6D). The primed-like flat colony morphology of GKO (Fig. 5A) was obviously reverted by simultaneous KO of Ampk (i.e., DKO) (Fig. 6F), which was consistent to the levels of primed pluripotency markers (Fig. 6G).

**Figure 6.**
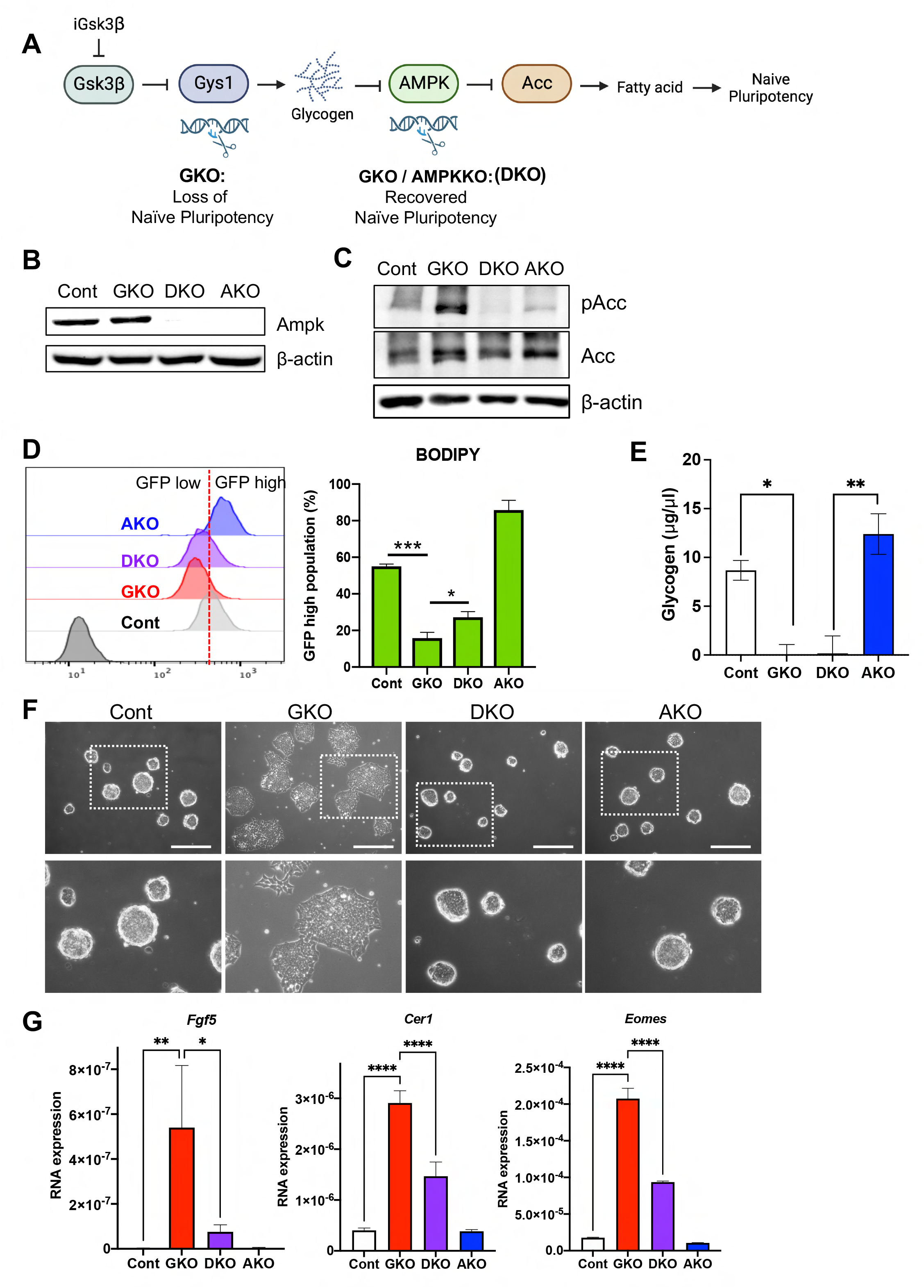
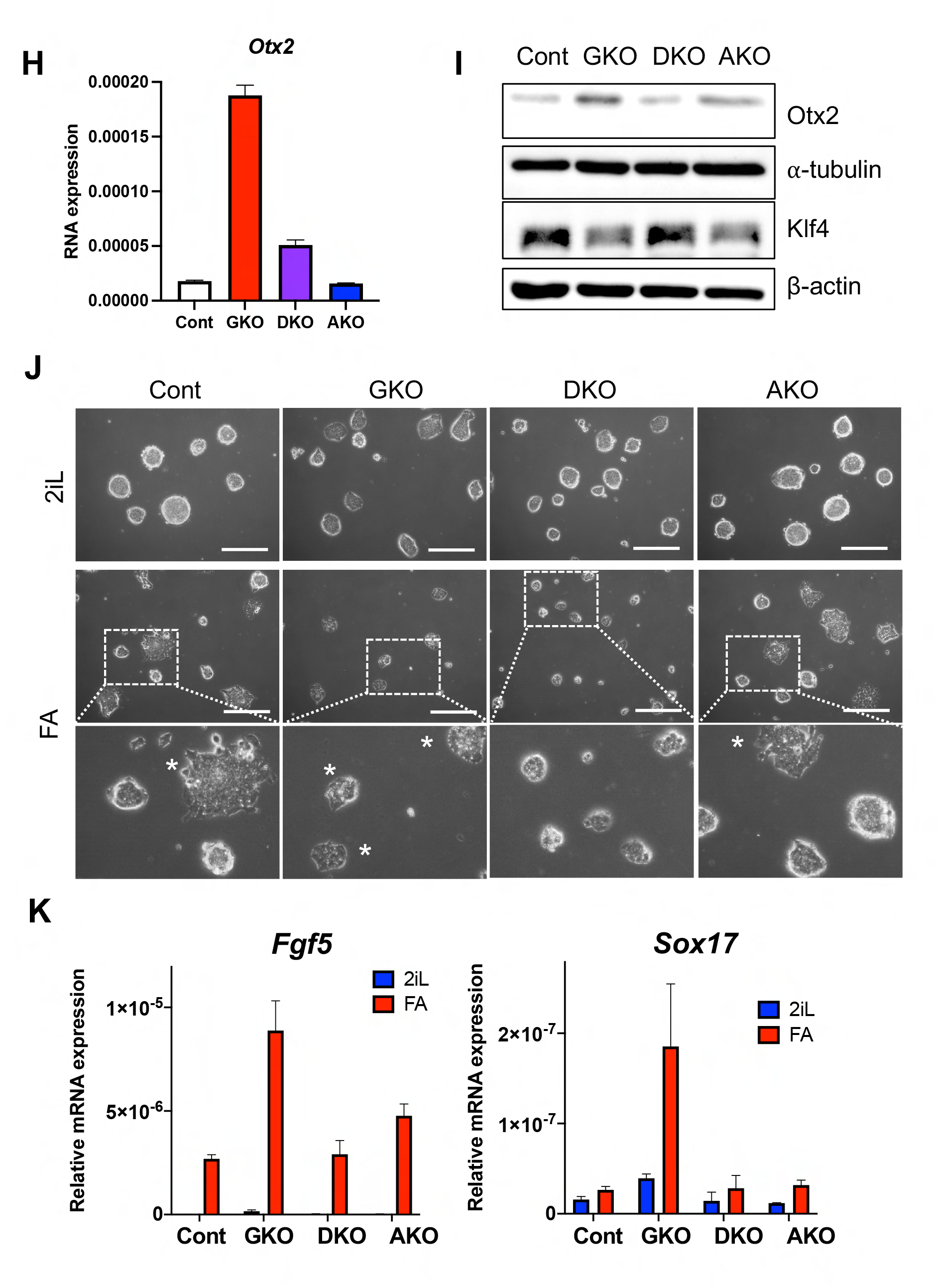
(A) Graphical scheme of GKO, AKO and DKO. (B) Immunoblotting analysis for indicative proteins (Ampk in Cont, GKO, DKO and AKO, *β*-actin was used as a loading control). (C) Immunoblotting analysis for indicative proteins (pACC, ACC in Mock, GKO, DKO and AKO, *β*-actin was used as a loading control). (D) Flow cytometry of BODIPY 493/503 staining in Cont, GKO, DKO and AKO (left panel), quantification of the populations from the flow cytometry (right panel). (***, p < 0.001, *, p < 0.05, n = 3). (E) Intracellular glycogen level of Cont, GKO, DKO and AKO. Glycogen assay kit was used. (F) Brightfield images of Cont, GKO, DKO, AKO (scale bars = 200μm). (G) Relative mRNA expressions for *Fgf5, Cer1, Eomes* in Cont, GKO, DKO and AKO under LIF+2i media condition. (H) Relative mRNA expression for *Otx2* in Cont, GKO, DKO and AKO. (I) Immunoblotting analysis for indicative proteins (Otx2, Klf4 in Cont, GKO, DKO and AKO), *β*-actin and ⍺-tubulin were used as loading controls. (J) Brightfield images of Cont, GKO, DKO, AKO under indicative media conditions. Primed conversion was performed by adding KOSR, bFGF and Activin A for 48 hours. (K) Relative mRNA expressions of *Fgf5* and *Sox17* under indicative media conditions. Primed conversion was performed by adding KOSR, bFGF and Activin A for 48 hours. Cont: Control, GKO: *Gys1* KO, AKO: *Prkaa1* KO, DKO: *Gys1* KO and *Prkaa1* KO, KOSR: Knock Out Serum Replacement, 2iL: 2i + LIF, FA: bFGF + Activin A.

The hypothesis that timely Ampk activation due to progressive loss of glycogen at the blastocyst stage (i.e., naïve ESCs) triggers the signal to epiblast transition (i.e., primed ESCs) was also supported by the significant upregulation of Otx2, one of the intrinsic determinants of epiblast transition ^45^, both at the transcription (Fig. 6H) and protein level (Fig. 6I and Extended data Fig. 5B), along with an inverse correlation with Klf4, a solely sufficient factor to induce the transition from epiblast stem cells to naïve cells ^46^. In turn, these ESCs were next induced to transition to a primed state with Fgf2/Activin A. Upon 48 hours of exposure to Fgf2/Activin A, a flat-like colony morphology was dominant in GKO, whereas both typical flat-like and dome-shaped colony morphologies were observed in the control (Fig 6J). Intriguingly, DKO maintained a dome-shaped colony morphology (Fig. 6J) upon primed induction. The results determined by colony morphology were confirmed by the expression levels of typical primed pluripotency markers (e.g., *Fgf5* and *Sox17* ^33^) (Fig. 6K). Taken together, these results suggest that activation of Ampk following the reduction of glycogen in naïve ESCs primes the transition to primed states.

## Discussion

Due to the limited number of *in vitro* models of mammalian embryos, the biological characterization of pre- and post-implantation embryos has remained challenging. Since the establishment of novel methods that allow for the *ex-vivo* production of blastoids (blastocyst-like structure) ^47^, extensive efforts have been made to establish artificial multi-cellular structures to mimic mouse and even human embryo development ^48^. Nevertheless, naïve and primed ESCs, which allow for extensive biochemical and molecular studies, provide valuable insights into the regulatory mechanisms underlying pluripotency in pre- and post-implantation embryos, respectively ^1, 49, 50^. The findings of this study highlight the importance of glycogen as a signaling molecule in the regulation of Ampk activation during early embryonic development. While glycogen has long been simply recognized as an energy reservoir, this study provides evidence that glycogen levels in the blastocysts can also serve as a regulatory mechanism to modulate Ampk activity. The observation that glycogen levels in mouse blastocysts decrease prior to implantation ^51^ suggests that timely Ampk activation through glycogen expenditure may occur *in vivo* during the pre-implantation stage of embryo development. Indeed, the critical role of Ampk activation during early embryogenesis is supported by genetic studies showing that knockout of Ampkα1/α2 results in embryonic lethality at approximately 10.5 days post-conception ^52^, as well as failure of differentiation of Ampkα/β KO ESCs ^53^. As shown in the cell model in this study (Fig. 3G), the sustained inhibition of Gsk3α/β by Netrin-1 signaling ^4^ would likely readily lead to Ampk activation when glycogen is depleted in the embryo under constant inhibition of Gsk3 activity through either Netrin-1 signaling in embryos or a chemical inhibitor of Gsk3β in naïve ESCs.

The role of fatty acid metabolism, specifically fatty acid β-oxidation (FAO), in reproduction biology and embryonic development has been extensively characterized ^54^. Key genes involved in fatty acid synthesis such as Fasn ^7^ and Acc ^8^ have been shown to be essential for early embryo development, which is consistent with our observation that chemical inhibition of Acc can induce cell death in naïve ESCs (Fig. 2H). Consistently, treatment with L-carnitine, which promotes FAO and potentially decreases fatty acid levels in embryos, has been shown to improve pregnancy rates following embryo transfer in bovine models ^55^ and humans ^56^. Furthermore, animals with higher levels of fatty acid in embryos such as cows, pigs, and cats ^10^ exhibit longer implantation times compared to mice, which have lower levels of fatty acid in embryos ^57^. This suggests that fatty acid metabolism, which is regulated by Ampk activity, may play a role in determining the timing of implantation in different species. We also examined the effect of GKO on teratoma. Teratomas derived from GKO were found to be significantly larger and comprised fewer undifferentiated cells compared to the controls (Fig. 5H and Extended data Fig. 3B). Notably, the variation of undifferentiated cells in GKO teratomas was significantly reduced compared to that of control teratomas (Extended data Fig. 4C), which would result from the synchronized differentiation of GKO due to lack of glycogen. Another question to be addressed is the molecular mechanism through which fatty acids delay the exit of naïve pluripotency. Previous studies have demonstrated that fatty acid metabolism, including FAO, provides acetyl carbon for histone acetylation ^58^ and is critical for providing a carbon source for pluripotency maintenance ^59^. This interesting hypothesis can be examined in follow-up studies.

Taken together, our results suggest that glycogen stored in blastocysts serves as a signaling molecule that controls timely Ampk activation, which in turn affects the level of fatty acids and transition to primed pluripotency. This could potentially explain the differences in fatty acid levels observed in embryos of different animal species, which have distinctly different implantation periods. These findings provide new insights into the role of glycogen as a signaling molecule in early embryonic development and shed light on the molecular mechanisms underlying the regulation of pluripotency and implantation.

## Availability of data and materials

The datasets used and/or analyzed during the current study are available from the corresponding author on reasonable request.

## Acknowledgement

This work was supported by a grant from the National Research Foundation of Korea (NRF-2020R1A2C2005914) and a grant from Regenerative Medicine funded by Ministry of Science and ICT, and Ministry of Health and Welfare (Grant number RS-2022-00070316), Republic of Korea.

