## Supplementary figures and images for "Ampk activation by glycogen expenditure primes the exit of naïve pluripotency"

### Extended Data Figures

**Extended Data Fig. 1**

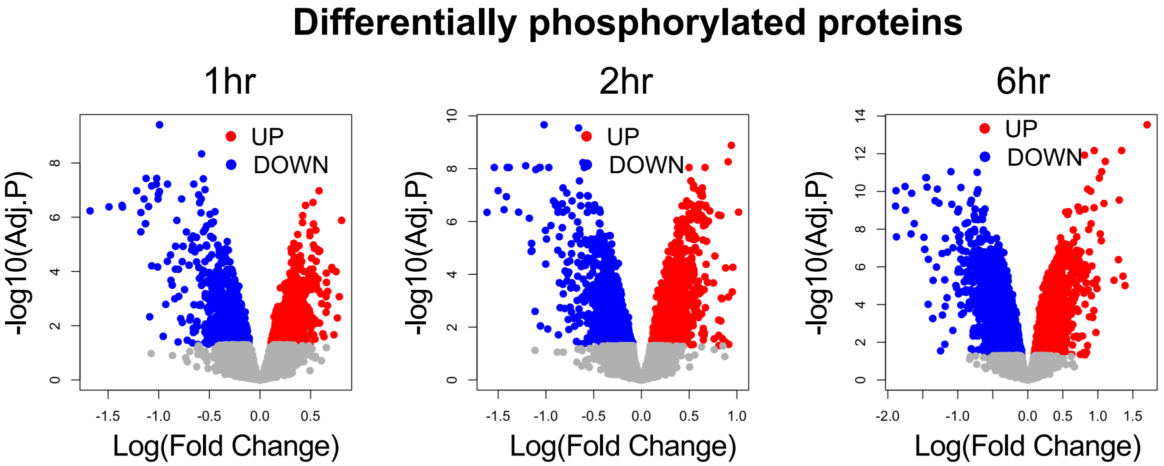

#### Extended Data Fig. 2

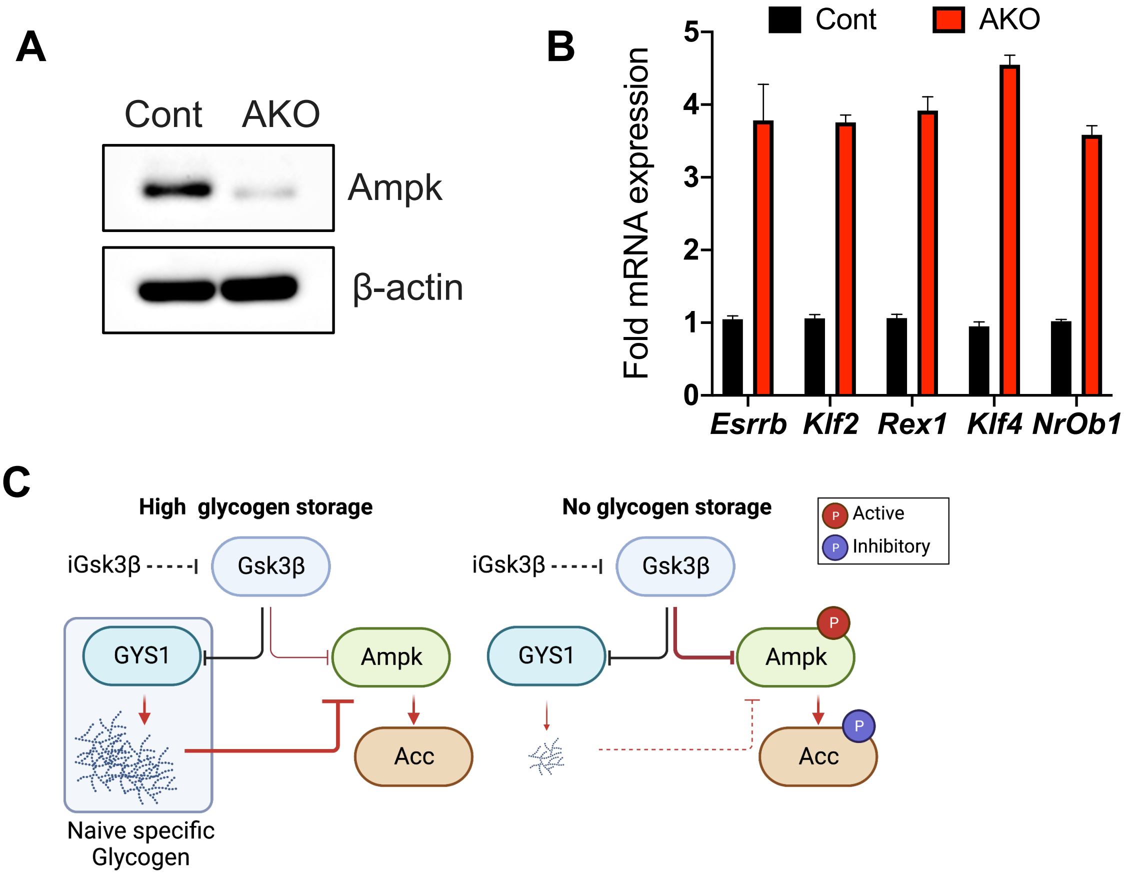

##### Extended Data Fig. 3

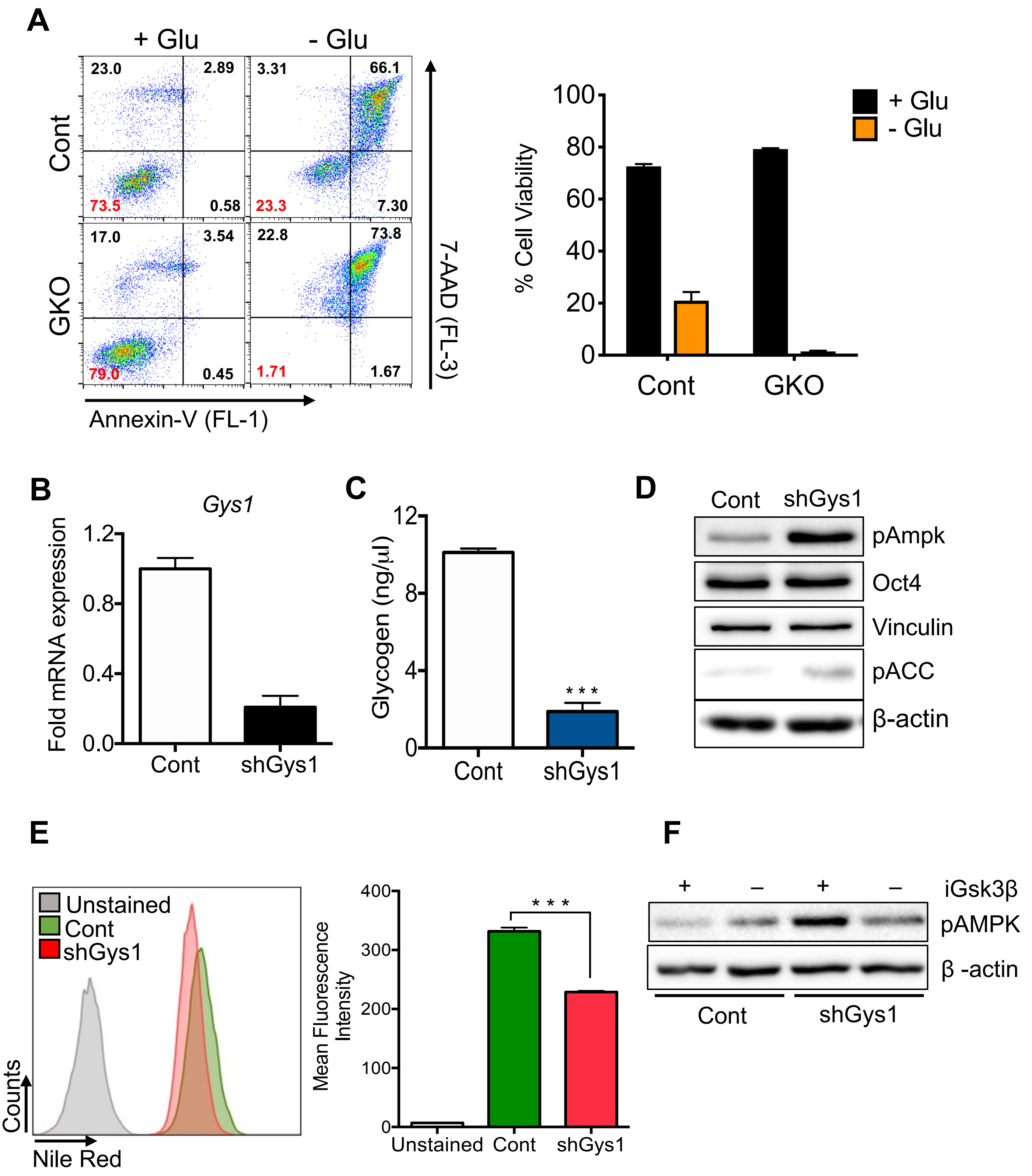

**Extended Data Fig. 4**

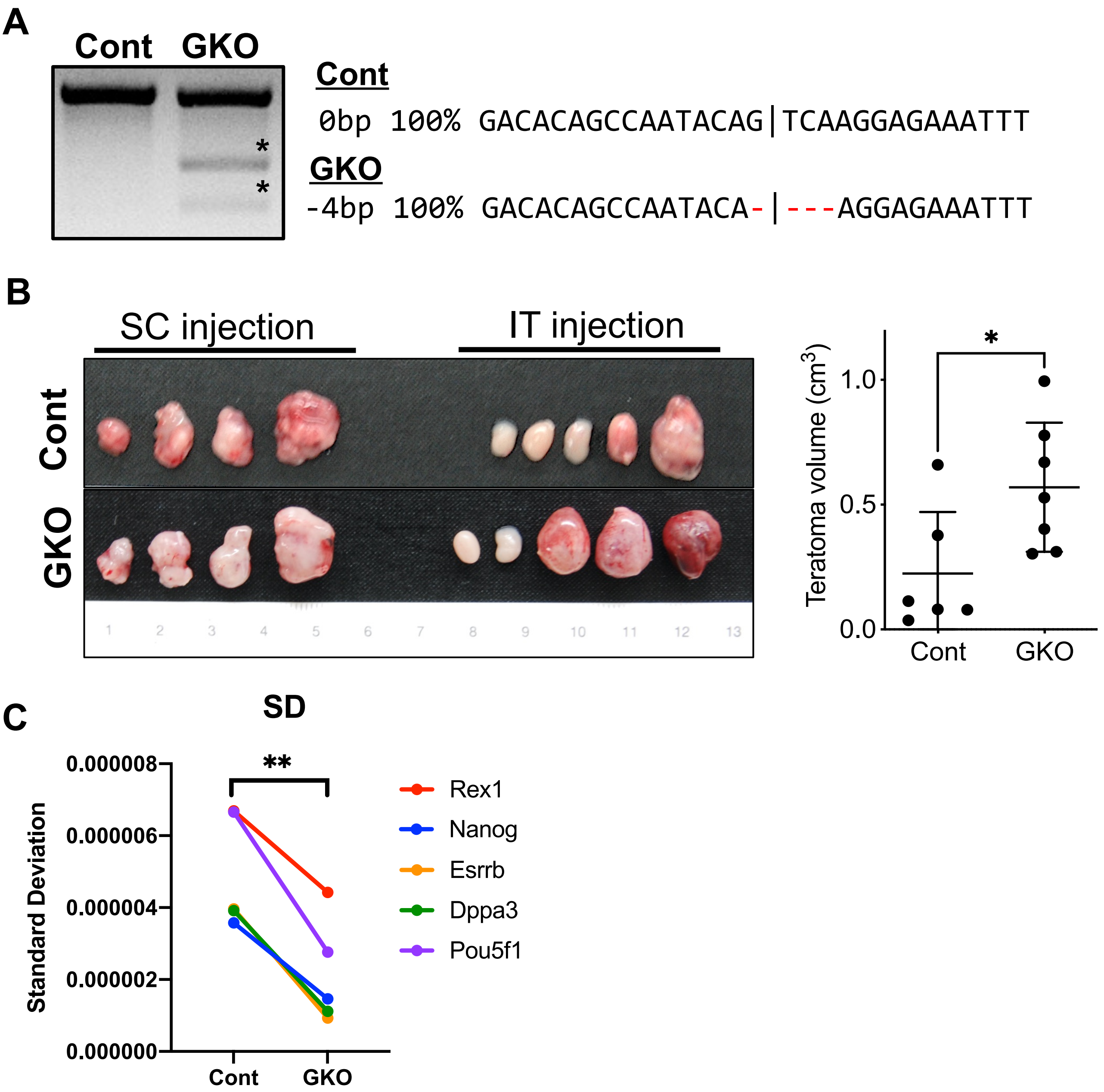

#### Extended Data Fig. 5

A

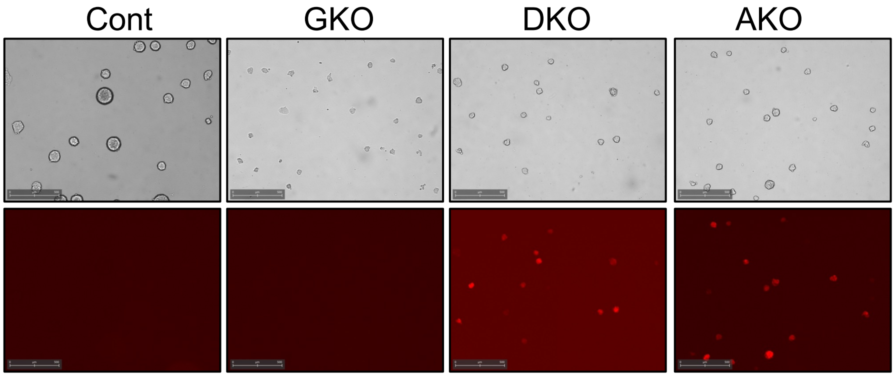


B

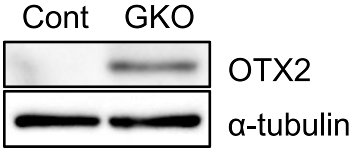
