## Extended Data Figure legends for "Ampk activation by glycogen expenditure primes the exit of naïve pluripotency"

### Extended data Fig. 1

Volcano plot of differentially phosphorylated proteins after 1,2 and 6hr of treatment compared to SL.

### Extended data Fig. 2

(A) Immunoblotting analysis for Ampkα, *β*-actin was used for loading control. (B) Fold mRNA expressions for *Esrrb*, *Klf2*, *Rex1*, *Klf4*, *NrOb1* in Cont and AKO.

### Extended data Fig. 3

(A) Flow cytometry for Annexin-V and 7-AAD staining of Cont and GKO with [+Glu] or without [-Glu] Glucose (left panel), quantification of live cell population (rignt panel). (B) Fold mRNA expression of *Gys1* in before [Cont] and after transient knock down of Gys1 [shGys1]. (C) Intracellular glycogen level before [Cont] and after transient knock down of Gys1 [shGys1]. (D) Immunoblotting analysis for indicative proteins (pAmpkα, Oct4, Vinculin, pAcc, *β*-actin was used for loading control) before [Cont] and after transient knock down of Gys1 [shGys1]. (E) Flow Cytometry of Nile Red staining before [Cont] and after transient knock down of Gys1 [shGys1] (left panel), quantification of Mean Fluorescence Intensity (right panel). (F) Immunoblotting analysis for pAmpkα (*β*-actin was used for loading control) before [Cont] and after transient knock down of Gys1 [shGys1].

### Extended data Fig. 4

(A) T7E1 assay for WT and GKO in OG2$^{+/-}$GOF6$^{+/-}$ cell line (left), sequence information of targeted *Gys1* from WT and GKO in OG2$^{+/-}$GOF6$^{+/-}$ cell line (right). (B) Representative images of Teratoma generated from Cont and GKO (left), representative graph of teratoma volume generated from Cont and GKO (right). (C) Intra-group standard deviation of gene expression in control and GKO teratoma samples.

**Extended data Fig. 5**

(A) Brightfield and RFP images for Cont, GKO, DKO and AKO. (B) Immunoblotting analysis for indicative proteins (OTX2 in Cont and GKO, α-tubulin was used as a loading control).
